# Trans-homologous interactions identified in Hi-C data are associated with embryonic development

**DOI:** 10.1101/2025.04.25.648339

**Authors:** Magdalena A. Machnicka, Aleksander Jankowski

## Abstract

Interactions between chromosomes forming homologous pairs are known to be widespread in the fruit fly *Drosophila melanogaster* and present in other organisms. While these trans-homologous interactions affect regulation of gene expression, many details regarding their location, function and mechanisms of action are still unknown. The increased availability of phased Hi-C datasets made it possible to reliably infer such trans-homologous interactions.

We have developed a tool called TransContactHiC to detect trans-homologous chromatin interactions in phased Hi-C data and we used it to re-analyze published in situ Hi-C data for heterozygotic *Drosophila melanogaster* cells: embryos and a fully differentiated cell line. We further assessed the frequency of cis- and trans-homologous interactions for equally sized genomic bins, and detected triads of genomic elements which exhibit significantly different frequency of trans-homologous interactions compared to the cis-homologous counterpart.

We show that triads of genomic elements exhibiting significantly higher frequency of trans homologous interactions are characterized by their clustering within broader regulatory regions, proximity to DNase-seq peaks and presence of disrupted transcription factor motifs at the affected locus. We also show examples of annotated active *cis*-regulatory modules which might be affected by these differential trans-homologous interactions. Overall, our results show the functional relevance of the detected triads.

## INTRODUCTION

Three-dimensional organization of chromatin into compartments and domains together with close physical interactions (known also as chromatin contacts) between genomic loci are essential for gene expression regulation (Hafner & Boettiger, 2023; Delaneau *et al*, 2019; Xu *et al*, 2020). Most often the interacting loci reside on the same chromosome (forming intrachromosomal interactions) but they can also be located on different chromosomes (forming interchromosomal interactions). A special case of the interchromosomal interactions are interactions between two chromosomes from the homologous pair. They are called trans-homologous (transh) interactions, in contrast to cis-homologous (cish) interactions which link loci located on the same chromosome from the homologous pair. Trans-homologous interactions have been observed in several organisms, first in *Drosophila melanogaster* within the bithorax complex (Lewis, 1954).

Significant homolog proximity has been observed in diploid budding yeast (*Saccharomyces cerevisiae*) (Kim *et al*, 2017), where the strength of interactions between homologs varies across both growth conditions and the genome. In mammals it is mainly observed during meiosis, but it also happens outside the germline where it is most often tightly regulated and occurs at specific loci (reviewed by (Apte & Meller, 2012)). However, more widespread homolog pairing during interphase has been recently reported in pig (*Sus scrofa*), where it positively correlated with the active A compartment and was suggested to be a source of additional layer of transcriptional regulation (Lin *et al*, 2024). However, an opposite phenomenon has been reported as well, with abnormal somatic pairing in human leading to disruption of gene expression and pathogenic processes (Koeman *et al*, 2008).

Highly structured genome-wide homolog pairing has also been extensively described in fruit flies (*Drosophila melanogaster*) (AlHaj Abed *et al*, 2019; Erceg *et al*, 2019). In this species, paired homologous chromosomes exhibit high interaction frequencies along their entire length, and transh interactions form structural features similar to cish interactions: interaction peaks, domains and compartments. The fly genome can be divided into tightly and loosely paired regions, and this division is associated with chromatin activity, location of compartments and gene expression.

The role of transh interactions and mechanisms through which they may influence gene expression regulation are only partially known. Regulatory interactions between homologous chromosomes, known as transvection, take place when transh interactions are imperative for proper gene expression through genetic complementation (Fukaya & Levine, 2017). The occurrence of transvection can be tested by inducing a disruption of transh interactions (e.g. by translocation) and confirming that this disruption leads to a change in phenotype. Transvection may rescue a regulatory interaction in case of dysfunction of one of the loci (Fukaya & Levine, 2017) or result in sex-biased expression (Galouzis & Prud’homme, 2021). In the fly embryos, transh interactions are involved in early developmental processes such as zygotic genome activation (Erceg *et al*, 2019). Even though homolog pairing is already present in very early stages of *Drosophila* development, it is more pronounced in differentiated cells, suggesting that the establishment of trans-homologous interactions takes place during cell differentiation (AlHaj Abed *et al*, 2019).

Transh interactions can be detected using the *in situ* Hi-C technique (Rao *et al*, 2014), which combines the original Hi-C protocol with nuclear ligation, ensuring that interactions under study are located within the nucleus of a single cell. When performed on heterozygous cells, this technique allows for identification of interactions between chromosomes which form homologous pairs. In this study, we present our tool, TransContactHiC, which we use to identify transh and cish interactions in the in situ Hi-C datasets from *Drosophila* hybrid cell line and embryos (AlHaj Abed *et al*, 2019; Erceg *et al*, 2019). Detailed characterization of the identified interactions provides further evidence for the involvement of transh interactions in regulation of gene expression.

## RESULTS

### Identification of trans-homologous interactions in *D. melanogaster* embryos and differentiated cells

We analyzed published in situ Hi-C data for heterozygotic *D. melanogaster* cells: whole embryos (WE) at 2-4h after egg laying (Erceg *et al*, 2019) and a fully differentiated cell line (Pat and Mat, PnM) (AlHaj Abed *et al*, 2019). The Hi-C reads were phased, which allowed us to identify and distinguish four types of interactions: cis-homologous (cish), trans-homologous (transh), cis-nonh (between non-homologous chromosomes within one haplotype) and trans-nonh (between non-homologous chromosomes representing different haplotypes).

We mapped the paired-end Hi-C sequencing data to the reference genome using bwa mem (Li, 2013), a local sequence aligner particularly suited for chimeric reads that needed to be split into more than one alignment. These alignments were further annotated using pairtools suite (Open2C *et al*, 2024) to detect and merge the alignments arising from the same DNA fragment being sequenced in both reads of the pair. Our tool, TransContactHiC, assigns haplotypes to alignments according to the underlying single nucleotide variants (SNVs) found in the read sequence (Figure 1A, see Methods). We further considered only chromatin contacts (interactions) where both alignments were phased.

**Figure 1.**
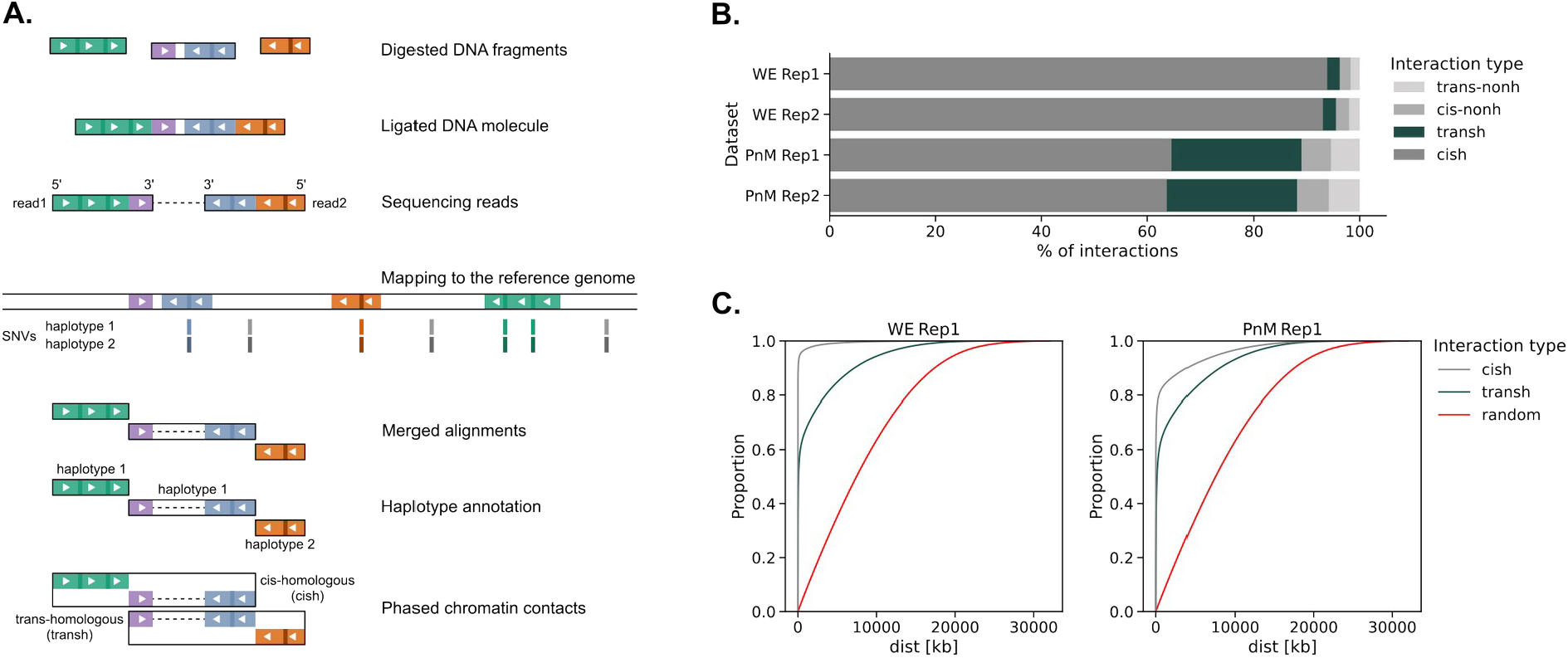
Identification of trans-homologous interactions in Hi-C data from *D. melanogaster*. **A**. Top to down: identification of cis-homologous and trans-homologous interactions by TransContactHiC. Genomic DNA is digested and re-ligated in a Hi-C experiment, and the resulting paired sequencing reads are mapped to the reference genome. The resulting alignments are merged when indicative of arising from the same DNA fragment. Haplotypes are assigned according to the underlying single nucleotide variants (SNVs) found in the read sequence. **B**. Percentage of interactions assigned to different types (cish, transh, cis-nonh and trans-nonh) identified in the fly embryos (WE) and Pat and Mat cell line (PnM). **C**. Empirical cumulative distribution functions (ECDF) plots for the distributions of distances between loci forming different types of interactions.

In total, we identified 33.1 million contacts in replicate 1 of the PnM data, 34.1 million contacts in replicate 2, as well as 24.1 and 24.5 million contacts respectively in replicate 1 and 2 of the WE data (Figure S1A). We found that transh interactions constitute a high percentage of all interactions identified in the PnM Hi-C data and that this percentage is approximately 10 times higher than in WE (24.4% *vs* 2.4%, Figure 1B). What is more, the absolute number of transh interactions is even more than 10 times higher in PnM than in WE (Figure S1A). For both WE and PnM, cish interactions are characterized by lower distances between the interacting loci than transh interactions (Figure 1C, Figure S1B). At the same time, the distances for both cish and transh interactions are lower than the distances between randomly selected pairs of genomic positions at the same chromosome.

### Genomic bin pairs enriched in transh or cish interactions

To identify regions of the genome in which transh interactions may have functional relevance, we tested for enrichment in transh interactions frequency with respect to cish interactions frequency. We wanted to detect cases in which transh interactions between the haplotypes are unexpectedly strong compared to cish interactions within one of the haplotypes. For this reason we have analyzed triads of genomic elements located at two interacting loci. Each element is defined by its genomic locus and haplotype: the first two are located on the first haplotype in the first and second locus, while the third is located on the second haplotype in one of these loci (Figure 2A). For each triad we consider two types of interactions: (i) cish interactions between the first two elements located on the first haplotype and (ii) transh interactions between the third element, located on the second haplotype, and one of the elements on the first haplotype.

**Figure 2.**
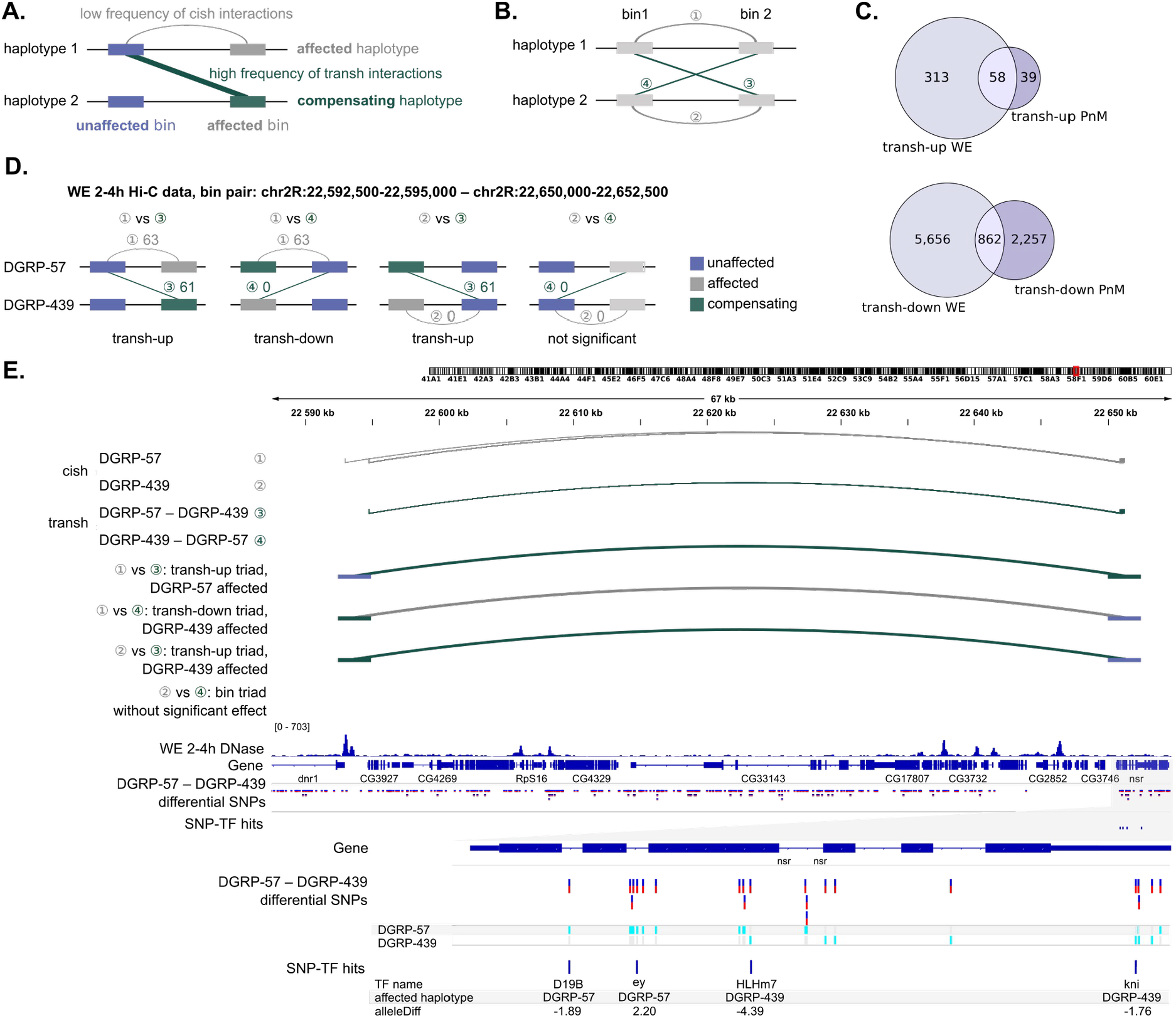
Identification of genomic element triads enriched in transh interactions. **A**. Schematic diagram of a transh-up bin triad with affected (depleted in chromatin contacts) and compensating (enriched in chromatin contacts) haplotypes marked. **B**. Schematic diagram of bin pair with all possible cish and transh types of interactions marked. **C**. Numbers of transh-up and transh-down triads identified in the fly embryos (WE) and Pat and Mat cell line (PnM). **D**.,**E**. Examples of bin triads enriched or depleted in transh interactions (transh-up or transh-down). **D**. All possible triads considered for this bin pair during the inference of transh interactions enrichment. **E**. Genome browser view presenting: cish and transh interactions (gray and green arcs), bin ranges, DNase-seq signal from embryos representing the same developmental stage (Reddington *et al*, 2020), SNVs differentiating the two haplotypes and SNVs potentially interfering with TF binding (identified with motifbreakR).

If the frequency of the transh interactions is higher than expected given the frequency of cish interactions, we call it “transh-up”. We use the names “unaffected” for the element involved in both cish and transh, “affected” for the element involved only in the unexpectedly weak cish interactions, and “compensating” for the element involved only in the unexpectedly strong transh interactions. Conversely, when the triad is characterized by lower than expected frequency of transh interactions we call it “transh-down”, and use the terms “affected” for the element having unexpectedly weak transh interactions, and “compensating” for the element having unexpectedly strong cish interactions.

To identify the transh-up and transh-down triads, we divided the genome into equally sized genomic bins (2.5 kb size) and used the counts of transh and cish interactions from two replicates as input for the DESeq2 tool (Love *et al*, 2014) (see Methods). For each pair of bins we tested four possible element triads for the difference between cish and transh interaction numbers (Figure 2B and D). In WE we were able to detect 371 triads in which the number of transh interactions was significantly higher than the number of corresponding cish interactions (transh-up). These triads represented 209 bin pairs. The number of transh-down triads identified in WE was approximately 18 times higher. In the same vein, in PnM we identified 97 transh-up triads for 70 bin pairs and approximately 32 times more transh-down triads (Figure 2C). An example of two transh-up triads and one transh-down triad within one pair of bins is shown in Figure 2E. In this case the number of transh contacts between the DGRP-57 haplotype in bin1 (chr2R:22,592,500-22,595,000) and DGRP-439 haplotype in bin2 (chr2R:22,650,000-22,652,500), equal 61 (counting both replicates together), is higher than expected given the number of cish contacts on haplotype DGRP-57 (63) and DGRP-439 (0) (two transh-up triads). At the same time, the number of transh contacts between DGRP-439 in bin1 and DGRP-57 in bin2, equal 0, is unexpectedly low compared to 63 contacts in cis on DGRP-57 (transh-down triad).

We noted that while PnM data are characterized by a much higher amount of transh contacts in general (both as a fraction and in absolute numbers) than WE, we called way less differential (transh-up or transh-down) triads for PnM than WE. What is more, the majority of transh-up triads identified in PnM are also transh-up in WE (Figure 2C). This is in line with the observations made by (AlHaj Abed *et al*, 2019) that upon cell differentiation the pattern of the transh interactions becomes more and more similar to the one of the cish interactions. This also suggests that transh-up genomic regions in PnM are infrequent and probably they are expected to have more functional relevance in WE.

### Transh interactions in transh-up bin triads are more clustered

Imbalance between transh and cish interactions in some genomic regions might be caused by local features of the haplotypes making one of them more predetermined for the interaction than the other. In such cases, we would expect the transh contacts to be clustered within the functionally relevant area. Indeed we could see that in transh-up bin triads, the individual transh contacts tend to be tightly clustered compared to cish contacts, both in transh-up and transh-down bin triads (Figure 3A and B). To quantify this clustering, we considered all bins from all the bin triads, taking transh-up and transh-down triads separately, and calculated the standard deviations of the positions of transh chromatin contacts and the cish chromatin contacts. We found a statistically significant difference in the distributions of these standard deviations for transh-up bin triads (Figure 3C and Supplementary Fig3A, see Methods). Such a significant difference has not been observed for transh-down comparisons. These results suggest that the enrichment in transh interactions observed for some of the bin pairs may point to functional differences between the haplotypes within these bins.

**Figure 3.**
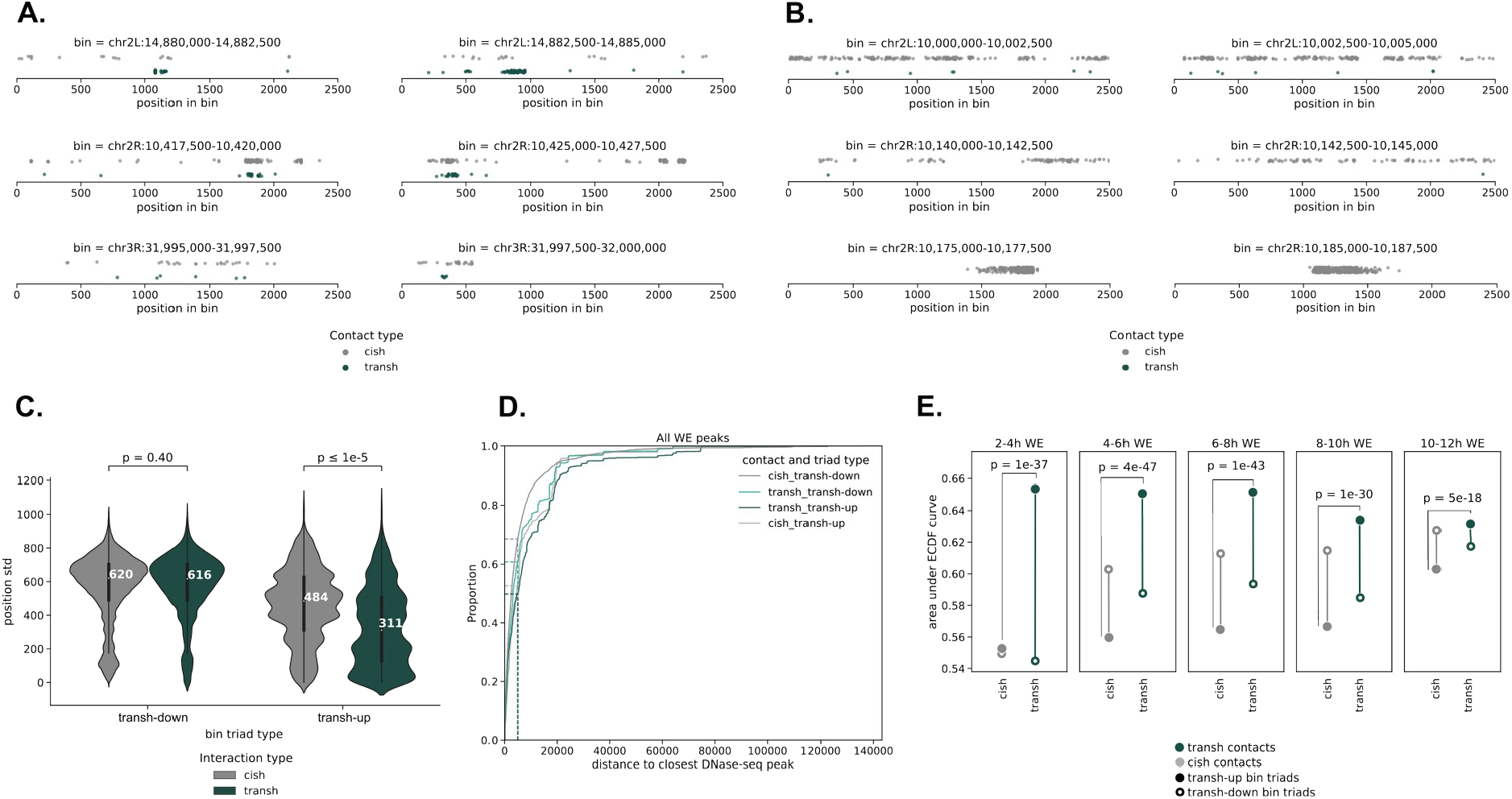
Transh contacts in transh-up bin triads are clustered and located close to embryonic DNase-seq peaks. **A**. Examples of bin pairs containing transh-up triads. Bins forming an interacting pair are plotted beside each other. Gray dots: positions of cish contacts, green dots: positions of transh contacts. **B**. Examples of bin pairs containing transh-down triads; colors used as in A. **C**. Distributions of standard deviations calculated for positions of contacts within bins from transh-up and transh-down triads identified in WE data. Median values are shown in white. P-values from two-sided Mann-Whitney *U* test. **D**. ECDF curves for distances between contact positions and closest DNase-seq peaks. All DNase-seq peaks from all tissues from (Reddington *et al*, 2020) were considered. Dashed lines indicate percentages of different contact type positions which have the closest peak within 5 kb. **E**. Dot plot with areas under the ECDF curves for distances between the closest DNase-seq peak and contact positions of different types. Only peaks from stages of embryonic development indicated above the plots were considered and only distances within 5 kb were included. The y-axis values were scaled from 0 to 1. P-values on the plot were obtained with the two-sided Mann-Whitney U test applied to the distributions of distances between transh and cish contact positions from transh-up and transh-down bin triads and their closest DNase-seq peaks. Only contact positions from the affected bins have been considered (see Figure S3B).

### Transh contacts in transh-up bin triads are located close to DNase-seq peaks

Functional chromatin contacts are often associated with gene regulatory events which require local openness of the chromatin (Lee *et al*, 2004; Shu *et al*, 2011). To further examine possible functionality of transh interactions in the identified transh-up bin triads, we assessed the distance between the individual chromatin contacts and closest DNase-seq peaks using embryonic DNase-seq data from (Reddington *et al*, 2020), focusing on contact positions located in the affected bins on both affected and compensating haplotypes (see Figure 2A and Figure S3B). Looking in the context of DNase-seq peaks from whole embryos at all developmental stages, we found that among all analyzed contact types the transh contacts from transh-up triads are characterized by the lowest proportion of contacts having the closest DNase-seq peak not further than 5 kb away (50% vs 53% for cish contacts from transh-up triads and 61-69% for contacts from transh-down triads, Figure 3D). The threshold of 5 kb is the value at which the short distance between the contact position and peak is interpretable in terms of regulatory activity. However, when DNase-seq peaks from whole embryos at different stages of development are considered separately, the transh contacts from transh-up triads appear to be located significantly closer to open chromatin than cish contacts from transh-up triads (within the range of 5 kb, p < 1e−17, Figure 3E). What is more, the highest disproportion between the distances is observed for the 2-4h whole embryo DNase-seq, which is the same time point and tissue as for the Hi-C data analyzed. This further supports the hypothesis about functional relevance of the transh-up triads identified in the WE dataset.

### Potential involvement of transh-up triads in regulatory processes

One of possible reasons for the regulatory impairment of one of the haplotypes is a genetic change hindering transcription factor (TF) binding. We used the motifbreakR tool (Coetzee *et al*, 2015), which assess the impact of small variants on TF binding sites, and focused on SNVs distinguishing the two haplotypes and located in transh-up triads. We checked if SNVs located in the haplotype which exhibits reduced contact frequency (affected) differ in their putative ability to change TF binding compared to SNVs located in the haplotype with increased contact frequency (compensating), and to SNVs located in the unaffected bins (Figure 2A).

MotifbreakR reports as hits SNV-TF pairs for which the SNV is expected to change the match between genomic sequence and DNA motif recognized by the TF. Each hit is assigned a value called alleleDiff which describes the change in the match: positive change indicates better match, negative – worse match. The higher the absolute value, the bigger the expected effect size. To capture SNVs which most likely have impact on regulatory processes we selected only those located within ±500 bp windows around summits of DNase-seq peaks. We observed that for all developmental time points, the affected haplotypes are characterized by the highest number of SNV-TF hits in which the alleleDiff is negative, while the compensating haplotypes have the lowest number of such hits (Figure 4A). The effect size is significantly different between the affected and compensating haplotypes only for hits with positive alleleDiff; compensating haplotypes tend to carry SNVs with higher positive impact on TF motif match than affected (Figure 4B, p = 0.002, merged DNase-seq windows from all time points). The median value of negative alleleDiffs is lower for affected haplotypes than for compensating, but the difference between the distributions is not significant (Figure 4B, p = 0.213).

**Figure 4.**
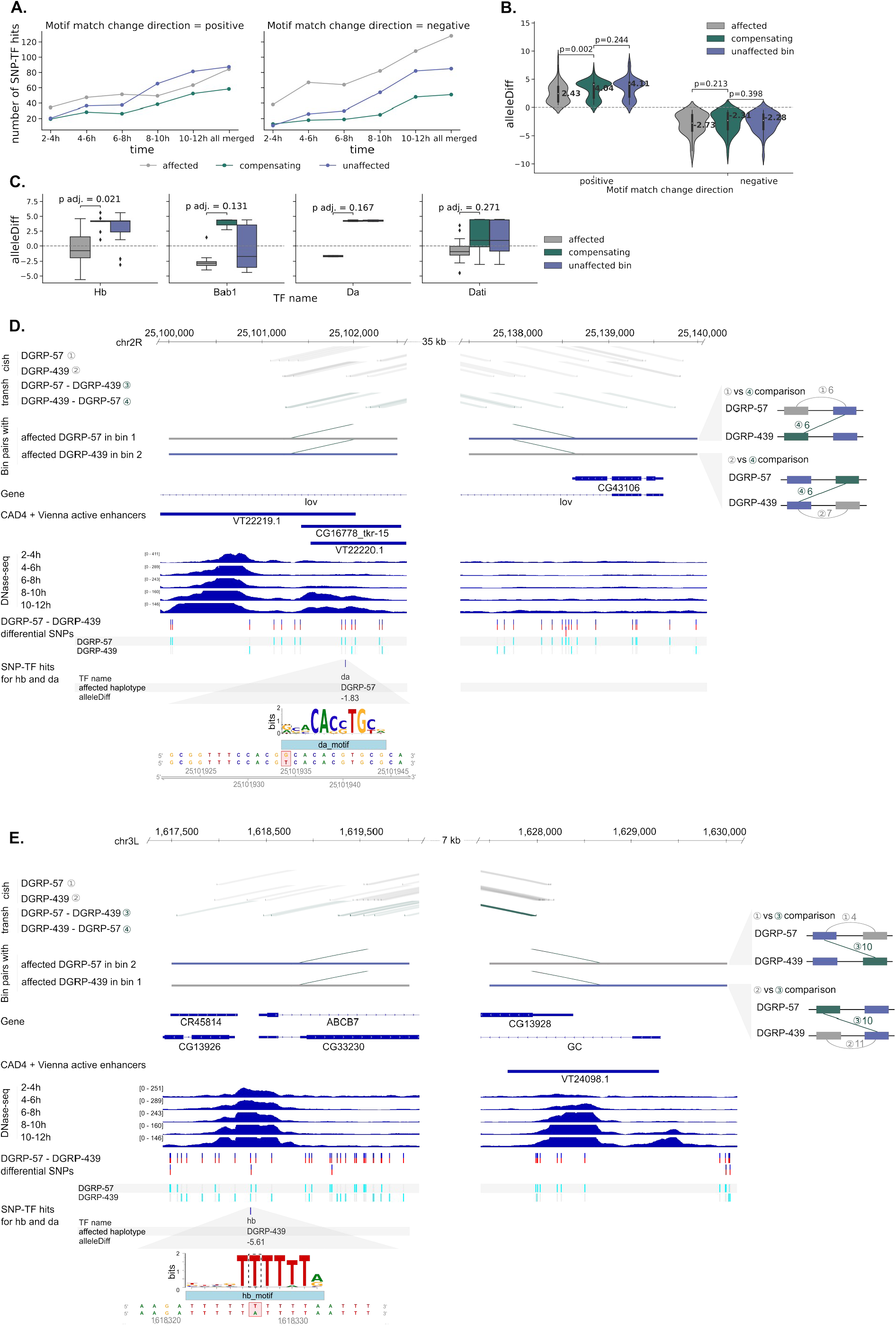
Transh-up bin pairs contain SNVs which might be involved in regulatory processes. **A**. Numbers of SNV-TF hits with positive or negative effect on motif match in affected or compensating haplotypes or unaffected bins. Only SNVs located within ±500bp windows around summits of DNase-seq peaks were included; x-axis: time points for the DNase-seq data **B**. Distributions of alleleDiff values for SNV-TF hits located in affected or compensating haplotypes or unaffected bins; p-values from the two-sided Mann-Whitney *U* test. **C**. Distributions of alleleDiff values in affected and compensating haplotypes or unaffected bins for the four selected TFs; alleleDiff – the difference between the motif match score on the reference allele and the score on the alternate allele calculated by the motifbreakR tool; padj. – p-values from the two-sided Mann-Whitney *U* test, Benjamini-Hochberg adjusted. **D and E**. Genome browser view for the chr2R:25101935G>T SNV influencing the Da motif instance (D) and chr3L:1618327T>A SNV influencing the Hb motif instance (E). Bins forming the pair under consideration and cish and transh interactions between these bins are shown. Active enhancers from the enhancer database (CAD4) described in (Cusanovich *et al*, 2018), available in Supplementary Table 13 therein, converted to dm6 genome assembly using liftOver (Hinrichs *et al*, 2006); DNase-seq signal from (Reddington *et al*, 2020).

We also looked for changes in binding sites of individual TFs and identified four which are characterized by negative alleleDiffs in affected haplotypes and positive in compensating haplotypes: Hb (Hunchback), Bab1 (Bric à brac 1), Da (Daughterless) and Dati (Datilografo). For all of them, the Mann-Whitney *U* test for the difference in the distributions of alleleDiffs in affected and compensating haplotypes gave p-values below 0.05, but only for Hb the p-value adjusted by the Benjamini-Hochberg procedure remained significant (padj. = 0.021, Figure 4C). Among them, Hb and Da are highly expressed during whole embryonic development, while expression of Bab1 and Dati is weaker and limited to selected, later stages in embryonic development, according to modENCODE RNA-seq data (Brown *et al*, 2014).

In Figure 4D,E (and Figure S4A,B) we present examples of sites in which a SNV in affected haplotype causes negative change in match between genomic sequence and Da or Hb motif. In the first of these examples, a SNV chr2R:25101935G>T, present in the DGRP-57 haplotype, modifies a putative Da binding site, located in an enhancer within an intron of the *lov* gene. This modification might lead to impaired regulation of the *lov* gene promoter via the enhancer located on the DGRP-57 haplotype, which could be compensated by transh interactions between the promoter region on the DGRP-57 haplotype and enhancer on the DGRP-439 haplotype. This enhancer overlaps three *cis*-regulatory modules (CRMs) which have positive impact on expression: VT22219.1, CG16778_tkr-15 and VT22220.1. The VT22219.1 module is known to regulate the *lov* gene and is active in embryonic stage 11 and 13 (approximately 6-10 h after egg laying at 25°C (Calderon *et al*, 2022)).

For VT22220.1, the target gene is unknown, but it is also active in stages 11 and 13 and additionally in stage 9 (Kvon *et al*, 2014), and CG16778_tkr-15 regulates *lov* gene during stages 10-12 (Brody *et al*, 2012) (data accessed through the REDfly database). The *lov* gene encodes a transcription factor of the BTB/POZ domain family (Bjorum *et al*, 2013; Koenecke *et al*, 2016). It is expressed at all stages of embryonic development as well as in some larval and pupal stages, and in adult males (Brown *et al*, 2014; Bjorum *et al*, 2013). It has a regulatory role during midline cell development (Wheeler *et al*, 2006) and its expression patterns in embryos suggest role in global re-structuring of the embryo and determination of neural lineages within the head (Bjorum *et al*, 2013). In larvae it regulates endopolyploid growth in several different tissues (Zhou *et al*, 2016, 2020). Misexpression of the *lov* gene was reported to lead to defective responses to gravity (Armstrong *et al*, 2006), behavioral defects in adults and locomotor defects in larvae (Bjorum *et al*, 2013).

In the second example, the Hb motif instance is negatively modified by the chr3L:1618327T>A SNV within a promoter of the *ABCB7* gene on the DGRP-439 haplotype. This promoter interacts with a region of open chromatin, containing one annotated positive CRM: VT24098.1, which is active in developmental stages 9 and 13 but its target gene is not known (Kvon *et al*, 2014). As a result the SNV may promote transh interactions between the intact promoter on the DGRP-57 haplotype and the enhancer on the DGRP-439 haplotype. The *ABCB7* gene, a putative *Drosophila* ortholog of the human mitochondrial ISC exporter ABCB7 (Metzendorf *et al*, 2009) encodes an ATP-binding cassette (ABC) transporter and is expressed at all developmental stages (Brown *et al*, 2014). It should be noted, however, that in both cases cish interactions involving the affected haplotype are only weaker than expected, not completely removed.

## DISCUSSION

Structural features of the chromatin have a great impact on its functioning, mainly due to the regulatory role of physical interactions between distant genetic elements. While the interactions within one of the homologous chromosomes (cis-homologous, cish) are most studied and best understood, the interactions between homologs (trans-homologous, transh) have also been found to have important functions. Recent developments in chromatin capture techniques, such as phased Hi-C experiments, allow for more detailed characterization of transh interactions. In this work we have developed the TransContactHiC tool, designed to infer transh chromatin contacts from Hi-C sequencing data. By re-analyzing available Hi-C datasets from *D. melanogaster* embryos and a differentiated cell line (Erceg *et al*, 2019; AlHaj Abed *et al*, 2019) we have demonstrated the potential of transh contact identification using TransContactHiC.

We identified a substantial number of transh contacts in both datasets and noted that while PnM data are characterized by a much higher amount of transh contacts in general (both as a fraction and in absolute numbers) than WE, we called way less differential (transh-up or transh-down) triads for PnM than WE.

The functional relevance of transh contacts identified with TransContactHiC is further supported by their clustering within DNA sequence, proximity to DNase-seq peaks and presence of disrupted TF motifs instances within sites enriched in transh contacts and depleted in cish contacts (at the affected locus). These results are in line with association of tight pairing with active genomic regions observed by (AlHaj Abed *et al*, 2019). We have also shown examples of annotated active CRMs which might be affected by the transh contacts identified with TransContactHiC. These results contribute to explaining the functions of transh contacts in *D. melanogaster* embryonic development. Unfortunately, phased Hi-C data are not available for adult flies, thus we could not apply TransContactHiC on differentiated cells other than the PnM cell line.

A limitation of our study is the reliance on Hi-C datasets with high enough coverage and high enough density of SNVs differentiating the haplotypes. In case of DGRP-57 x DGRP-439 cross, the average density of informative SNVs was one per 220 bp. Low coverage and lack of SNVs allowing for read phasing may lead to a high level of false negative results. Differential transh triads that we were able to identify in the re-analyzed fly datasets had a low number of chromatin contacts contributing to their identification (only 18% had at least 100 contacts). As embryos develop and tissues differentiate, functional transh contacts could get harder to detect, as they would be present only in a subset of cells assayed.

A possible remedy would be to perform a targeted chromatin capture experiment, such as Capture-C (Downes *et al*, 2022) or T2C (Kolovos *et al*, 2014), focusing on promoters of certain genes of interest. This would allow to gather higher sequencing read coverage of putative pairs of loci, and in turn to identify differential transh triads with greater statistical power. As seen in Figure 2AB, our approach could be easily adapted to use contact counts derived from these experiments, as our model does not require a broader genomic background – haplotype-resolved chromatin contacts linking the two loci of interest would be sufficient. Such analysis could focus on regions flanking transvection-mediating insulators (Piwko *et al*, 2019), enabling a better understanding of trans-homologous insulator-insulator interactions and the proposed formation of trans-homologous topological domains or hubs (Lim *et al*, 2018).

## METHODS

### Calling SNVs in DGRP-57 and DGRP-439 haplotype

Whole-genome sequencing data from the *Drosophila melanogaster* Genetic Reference Panel (Mackay *et al*, 2012) downloaded from Short Read Archive (experiment SRX021296 for DGRP-57, SRX021244 for DGRP-439) were mapped to the dm6 genome (FlyBase version FB2020_05, dmel_r6.36) using bwa mem (Li, 2013) version 0.7.17. The resulting BAM files were sorted by genomic coordinates using samtools (Danecek *et al*, 2021) version 1.10, and duplicates were identified using picard MarkDuplicates from Picard (https://broadinstitute.github.io/picard/) version 2.26.4.

Small polymorphisms such as SNVs, MNPs (multi-nucleotide polymorphisms) and indels (insertions or deletions) were called using Bayesian haplotype-based variant detector FreeBayes (Garrison & Marth, 2012) version 1.3.5 on both samples simultaneously, with disabled population priors. The results were filtered by vcffilter from vcflib (Garrison *et al*, 2022) version 1.0.3 using filter QUAL > 20, i.e. retaining only the variants with estimated probability of not being polymorphic less than 0.01. We further used vcflib to normalize complex haplotype calls into pointwise SNPs and indels, and selected only biallelic SNPs.

To separate the haplotypes in Hi-C data, we took only the variants that were homozygous both in DGRP-57 and in DGRP-439, and differed between these two haplotypes. The final set of differential SNPs consisted of 623,703 SNVs. These variants were also used to prepare haplotype-specific genomes (alternate references) in FASTA format using vcf-consensus from vcftools (Danecek *et al*, 2011) version 0.1.16.

### Identification of cish and transh chromatin contacts

Hi-C sequencing data from DGRP-57 (maternal)/DGRP-439 (paternal) heterozygous cross were downloaded from Short Read Archive (experiments SRX4887861 and SRX4887862: two replicates for 2-4h embryos; experiments SRX4887863 and SRX4887864: two replicates for the ‘Pat and Mat’ cell line). The reads were mapped to the same genome as above, considering both reads from a read pair separately, using bwa mem version 0.7.17 with options -E50 -L0 -5. The resulting BAM files (separate for read1 and read2) were further processed using samtools version 1.10. After sorting the reads by their name, bitwise flags indicating read1 or read2 were set, and the separate files were merged using samtools merge, followed by filling in mate-related flags and coordinates by samtools fixmate. Finally, optical and PCR duplicates were removed using picard MarkDuplicates from Picard version 2.26.4.

The alignments of filtered paired reads were further annotated using pairtools parse from the pairtools suite (Open2C *et al*, 2024) version 1.0.2 with options --walks-policy all --add-columns pos5,pos3 --no-flip. These options ensured that the alignments were reported along with their full genomic coordinates, and with their sequencing order preserved. The annotations, saved in .pairs.gz format, allowed for merging the alignments arising from sequencing the same DNA fragment in both reads forming the paired-end read.

Our tool, TransContactHiC, processes chromatin contacts by simultaneously reading the BAM file and corresponding .pairs.gz file, annotating the haplotype of each alignment according to the underlying SNVs. An important technical detail is to distinguish between two types of alignments: BAM alignments (in BAM files) are located in either read1 or read2 of the Hi-C read pair, while pairtools alignment (in .pairs.gz files) span one or two BAM alignments, and in the latter case they merge BAM alignments located in both read1 and read2.

The input files of TransContactHiC are: BAM file (containing multiple BAM alignments for each read pair), .pairs.gz file (containing multiple pairs of pairtools alignments for each read pair) and a .tsv.gz file containing SNVs. The pairtools alignments are first matched with BAM alignments. Haplotypes are then assigned to each of the pairtools alignments based on the read sequence stored in BAM alignments and the provided SNVs. The haplotype is annotated as “unknown” in case of no informative SNVs, or “ambiguous” if the underlying SNVs give conflicting annotations. The main output is the .tsv.gz file containing one row for each chromatin contact (suitable alignment pair from the input .pairs.gz file), along with haplotype information of both pairtools alignments. Keeping the cish and transh chromatin contacts together facilitates further downstream analysis. A BAM file with the annotated BAM alignments can also be saved.

### Identification of bin pairs with differential frequency of transh interactions

DESeq2 version 1.30.1 (Love *et al*, 2014) was used to identify bin pairs characterized by significantly higher or lower frequency of transh interactions compared to cish interactions. Interactions of each type in each bin pair were counted for both replicates. Differential analysis with DESeq2 was run on pre-filtered data (rows with mean counts below 1 excluded). Significant results were selected based on padj. < 0.05 threshold.

### Clustering of transh interactions

The level of clustering of interactions within bins was assessed by calculating standard deviations for genomic coordinates (positions) of all contacts of each type (cish/transh) within each bin. The distributions of standard deviation values for cish and transh interactions located in transh-up or transh-down bin pairs were compared with the Mann-Whitney *U* test.

### Proximity to DNase-seq peaks

DNase-seq peaks from different stages of *D. melanogaster* embryonic development from (Reddington *et al*, 2020) (accession codes: E-MTAB-8881 and GSE101581) were used to analyze the distance between positions of chromatin contacts and closest DNase-seq peak (using the bedtools closest tool (Quinlan & Hall, 2010). For the distance calculation in Figure 3D, we took the union of all peaks in all time points (2-4h up to 10-12h) and tissues (whole embryo, neuroblasts, neurons, myogenic mesoderm, visceral mesoderm, not neurons nor myogenic mesoderm, myogenic mesoderm without visceral mesoderm). For Figure 3E, only the whole embryo (WE) peaks were considered. The distributions of distances for cish and transh interactions located in transh-up or transh-down bin pairs were compared with the Mann-Whitney *U* test. Only contact positions located in the affected bins have been considered (as shown in Figure S3B).

### Regulatory functions of SNVs

SNVs located in transh-up bin pairs and differentiating between the two haplotypes were used as input for the motifbreakR tool (Coetzee *et al*, 2015) version 2.4.0 which calculates the difference in TF motif match between the reference and alternative allele (alleleDiff). Negative value of alleleDiff indicates that the presence of the SNV disrupts a TF motif instance while positive suggests better match. The motifbreakR analysis was run using modified dm6 reference genomes: for SNVs located in one of the haplotypes the dm6 genome with SNVs from the other haplotype inserted was used as reference. Only SNVs located within ±500bp windows from summits of the WE DNase-seq peaks were considered.

All TF motifs available in MotifDb version 1.32.0 for *D. melanogaster* were used for this analysis. For TFs represented by several motif matrices, alleleDiff values obtained for different matrices were grouped together. In cases when a SNV had identical alleleDiff scores for more than one matrix describing the same TF and when the length of these matrices and offsets describing the location of the match were identical, then only one of them was kept (identical matrices may be present in MotifDb under different IDs). Subsequently, alleleDiff values distributions were compared with the Mann-Whitney *U* test, multiple hypothesis testing correction was performed with the Benjamini-Hochberg method.

We used FlyBase (release FB2024_02, released April 23, 2024) to find information on phenotypes/function/stocks/gene expression etc. (Öztürk-Çolak *et al*, 2024). Data about active CRMs were accessed through REDFly version 9.6.4 (Keränen *et al*, 2022).

## Code availability

The code used to run the above analysis is available on-line: https://github.com/ajank/TransContactHiC.

## Author’s contributions

**Conceptualization:** Magdalena A. Machnicka, Aleksander Jankowski

**Data Curation:** Magdalena A. Machnicka, Aleksander Jankowski

**Formal Analysis:** Magdalena A. Machnicka, Aleksander Jankowski

**Funding Acquisition:** Aleksander Jankowski

**Investigation:** Magdalena A. Machnicka, Aleksander Jankowski

**Methodology:** Magdalena A. Machnicka, Aleksander Jankowski

**Project Administration:** Aleksander Jankowski

**Resources:** Aleksander Jankowski

**Software:** Magdalena A. Machnicka, Aleksander Jankowski

**Supervision:** Aleksander Jankowski

**Validation:** Magdalena A. Machnicka, Aleksander Jankowski

**Visualization:** Magdalena A. Machnicka, Aleksander Jankowski

**Writing – Original Draft Preparation:** Magdalena A. Machnicka, Aleksander Jankowski

**Writing – Review & Editing:** Magdalena A. Machnicka, Aleksander Jankowski

## Acknowledgements

This work was supported by the Polish National Agency for Academic Exchange (Polish Returns 2019, to AJ).

## SUPPLEMENTARY FIGURES AND TABLES

**Figure S1.**
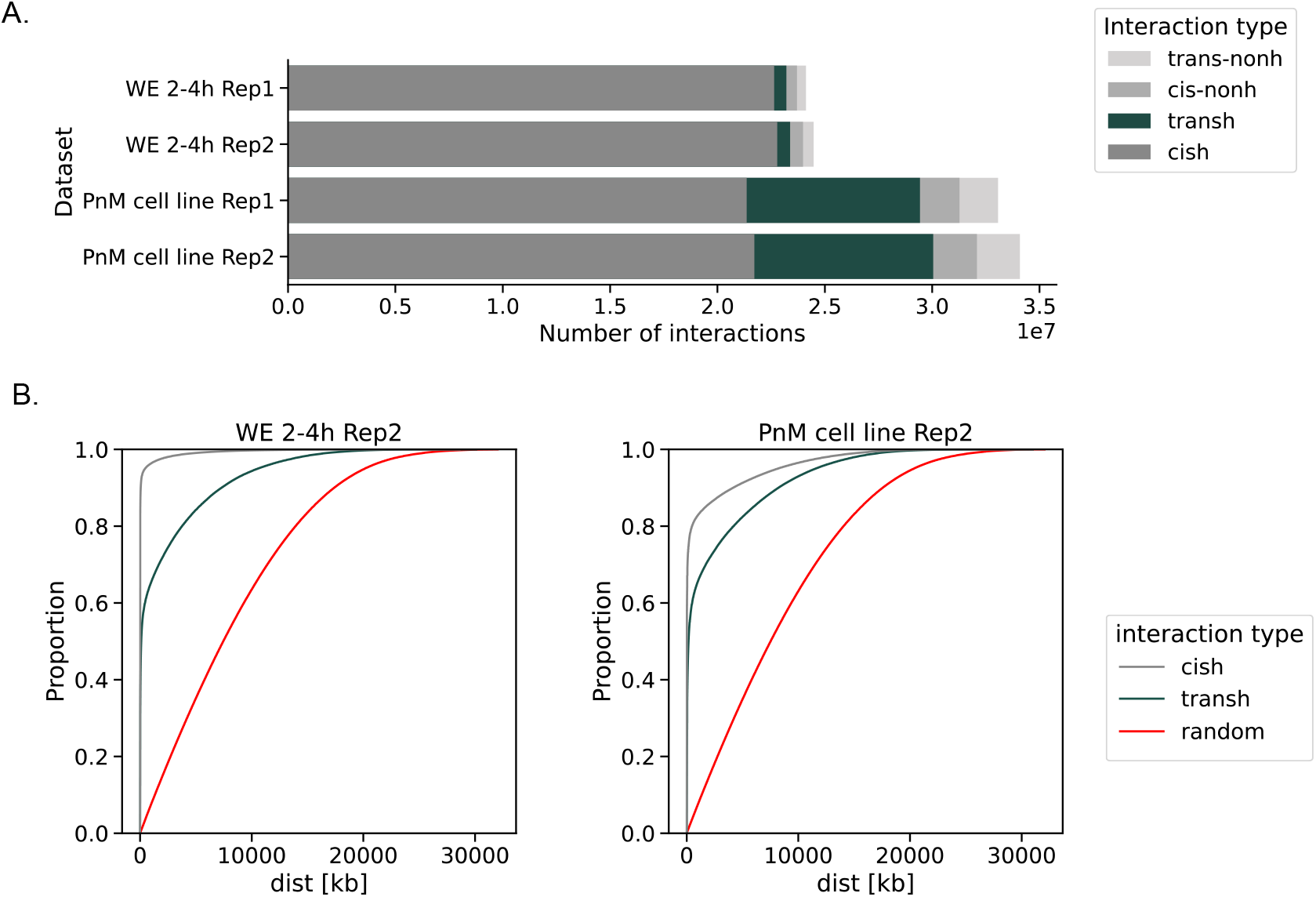
**A**. Absolute counts of different interaction types identified in the fly embryos (WE) and Pat and Mat cell line (PnM). **B**. ECDF plots for the distributions of distances between loci forming different types of interactions in Rep2.

**Figure S2.**
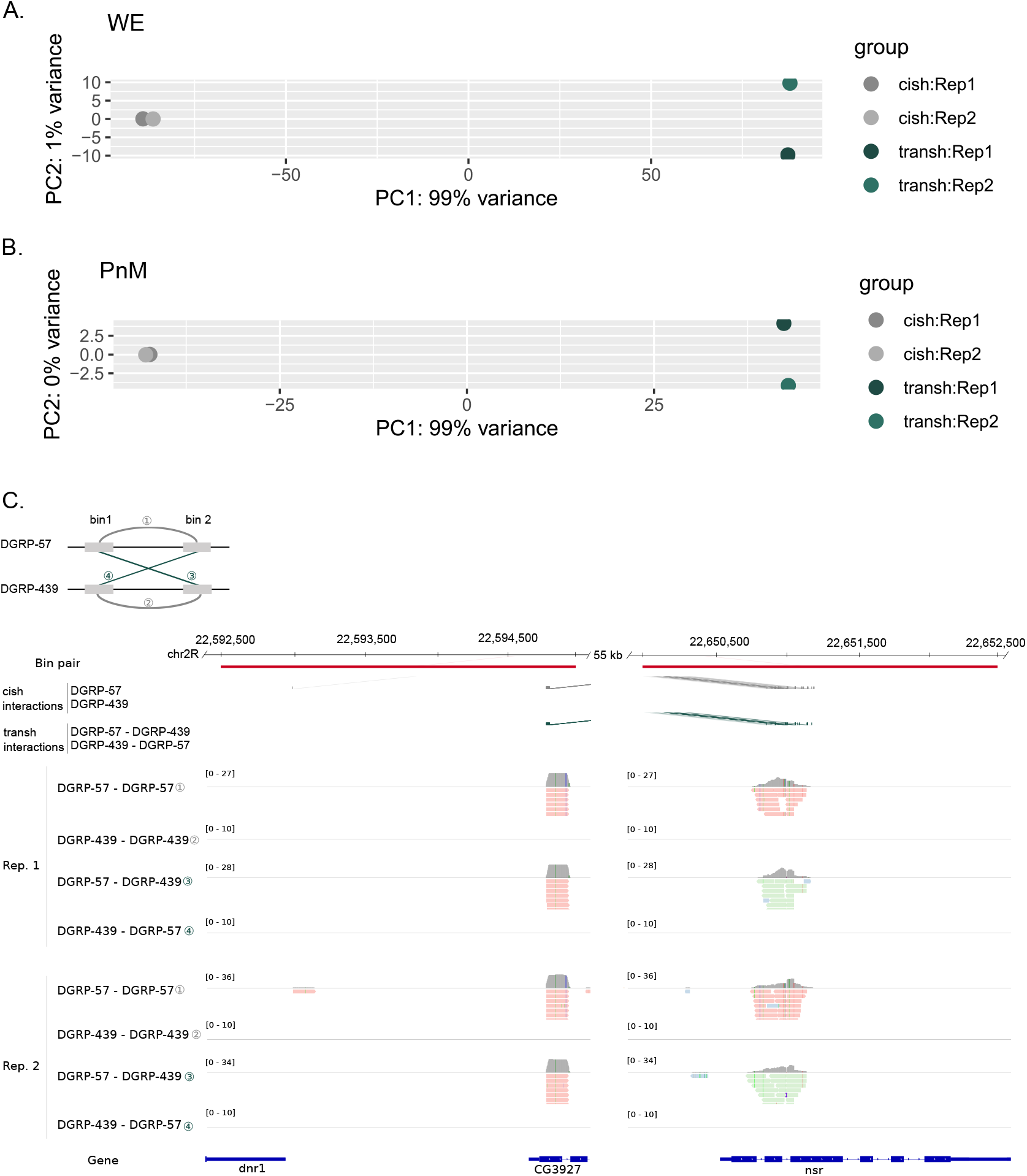
**A. and B**. PCA plots for counts of cish and transh interactions between bin pairs used for the identification of transh-up and transh-down comparisons with DESeq2. **C**. Genome browser view of the selected example of transh-up bin pair (associated with Figure 2DE). Tracks included in the presented view: location of the selected pair of genomic bins (in red), location of cish (in gray) and transh (in green) interactions linking bins from this pair, coverage and reads supporting each interaction type, presented separately for the two replicates. Reads are colored according to the haplotype they were assigned to (red – DGRP-57, green – DGRP-439).

**Figure S3.**
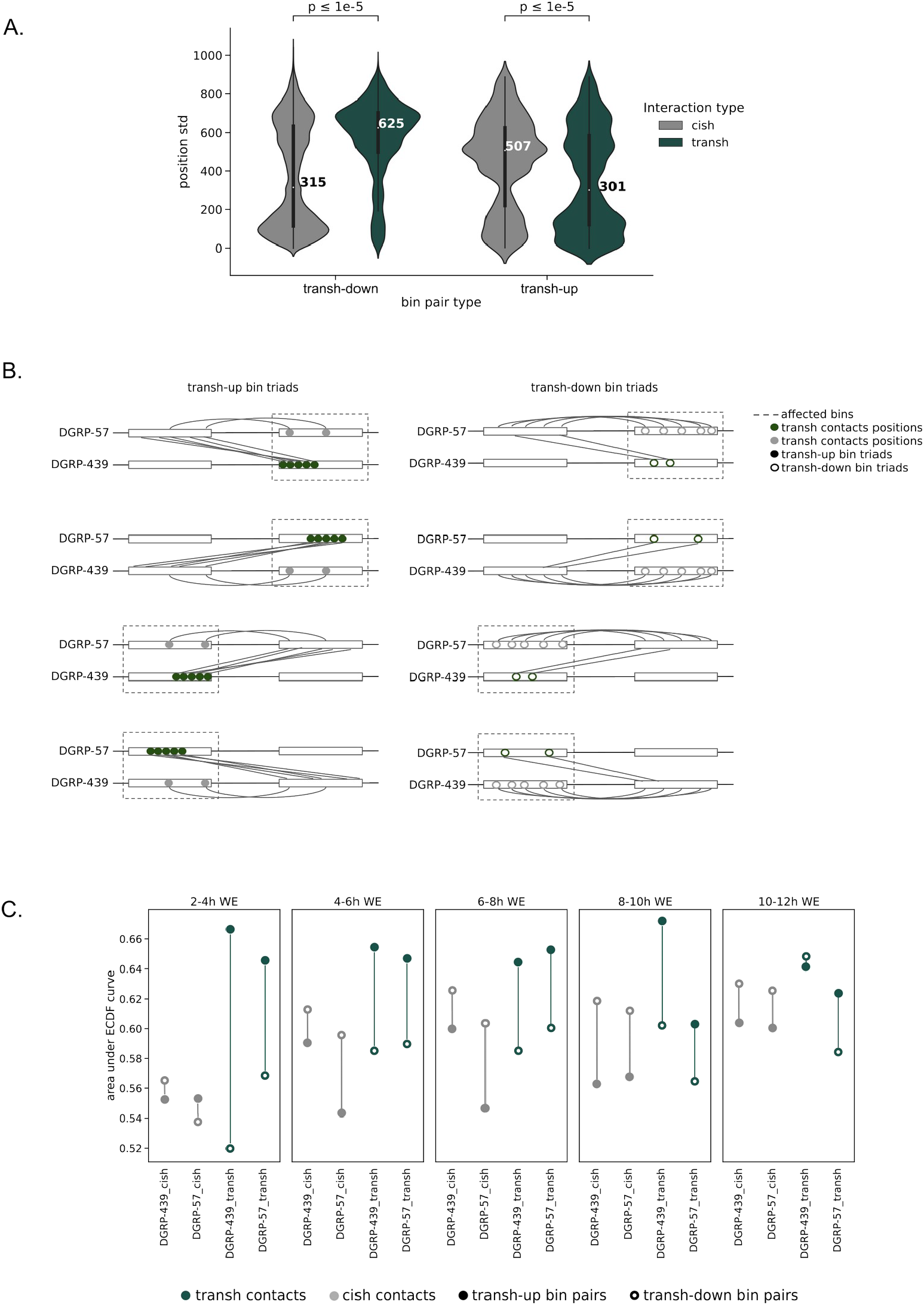
**A**. Distributions of standard deviations calculated for positions of contacts within bins from transh-up and transh-down bin triads identified in PnM data. Median values are shown. P-values from two-sided Mann-Whitney *U* test. **B**. Schematic diagram showing which contact positions have been considered for the analysis of distances between contacts and the closest DNase-seq peaks. **C**. Dot plot with areas under the ECDF curves for distances between the closest Dnase-seq peak (Reddington *et al*, 2020) and contact positions of different types. Contacts located on different haplotypes are shown separately. Only peaks from stages of embryonic development indicated above the plots were considered and only distances within 5 kb were included. The y-axis values were scaled from 0 to 1.

**Figure S4.**
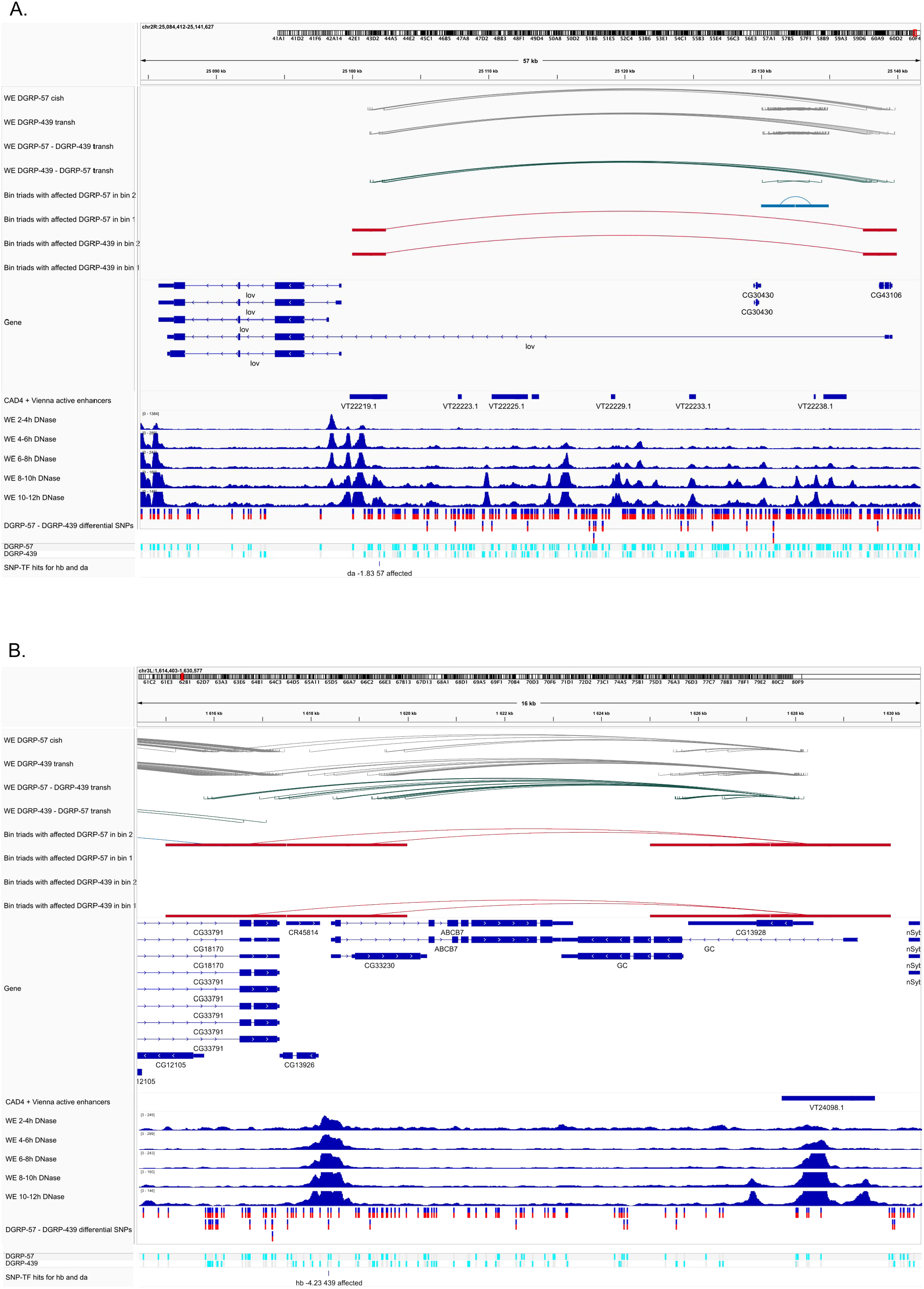
Additional genome browser views for the selected examples of SNVs disrupting TF motifs. **A**. chr2R:25101935G>T SNV influencing the Da motif instance, **B**. chr3L:1618327T>A SNV influencing the Hb motif instance.

**Table S1**. Coordinates of bin pairs with differential transh contacts in WE.

**Table S2**. Coordinates of bin pairs with differential transh contacts in PnM.

